# The algebra of community temperature indices

**DOI:** 10.64898/2026.08.02.742286

**Authors:** Matthew Spencer

## Abstract

The Community Temperature Index (CTI) was developed as a way to measure changes in a community over time in a way that reflects the temperature preferences of species. Several different forms of CTI are in widespread use, but their properties have not been systematically investigated. We argue that a CTI should preserve the algebraic structure of relative abundances given by the Aitchison geometry used in compositional data analysis. We show that this requirement leads to a new Aitchison CTI. The CTI has been used to classify species as warm- or cold-affinity, depending on whether an increase in their relative abundance increases or decreases the CTI. However, such classifications depend on relative abundances as well as on temperature preferences, which is undesirable. We show that the Aitchison CTI leads to a classification of species as relative warm- or cold-affinity that does not depend on relative abundances, with species’ contributions to changes in CTI that are consistent with the principles of population dynamics. We show that the Aitchison CTI is approximately linearly related to the most popular CTI in current use if relative abundances are approximately equal and temperature preferences for all species are close to their geometric mean. We provide a quantity analogous to an *R*^2^ for the Aitchison CTI, that tells us whether the CTI contains useful information. We illustrate our approach using data from a hard-substrate macrobenthos community.

## Introduction

The Community Temperature Index (CTI) was introduced by Devictor et al. (2008) as a way to measure changes in a community over time in response to environmental change. Devictor et al. (2008, p. 2744) defined the CTI as “the average of each individual’s STI present in the assemblage”, where the STI (Species Temperature Index) for a species is the mean temperature experienced by individuals of that species over its entire geographical range. No mathematical notation was used to make the original definition precise. However, the most common form of CTI (Supporting Information, Table S1 and Sections S1, S2, S3) appears to be as follows. Let *n* ∈ ℕ be the number of species in the community, and let the relative abundances of these species be **x** = (*x*_1_, *x*_2_, …, *x*_*n*_), where for each *i* ∈ {1, 2, …, *n*}, 0 *< x*_*i*_ *<* 1, and 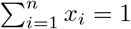. Then typically,

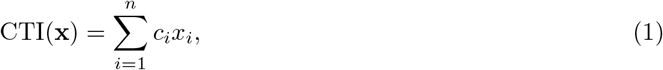

where *c*_*i*_ is the STI for the *i*th species (Gaüzère et al., 2019). Other forms of CTI are summarized in the Supporting Information (Section S2), although in many cases the form of CTI was described only in words, and was thus not entirely clear. Most papers using a CTI study changes in the CTI over time and/or space (Supporting Information, Table S1 and Section S3). In the rest of this paper, I refer to time throughout, but all the arguments also apply to space or some other variable. In addition, all the arguments apply to related indices in which the *c*_*i*_ are some constants other than STIs. There is also some interest in classifying species as warm- or cold-affinity, depending on whether an increase in their relative abundance increases or decreases the CTI (McLean et al., 2021), and then decomposing changes in CTI into changes in warm- and cold-affinity species. However, such classifications run into problems for the most common form of CTI, because whether a species is warm- or cold-affinity depends on the relative abundances of all species, not just on their STIs, requiring an arbitrary choice of reference CTI.

The typical choice of CTI (Equation 1) looks like a first-degree Scheffé polynomial in canonical form (Cornell, 2002, section 2.2). Such polynomials are widely used to model properties of a product such as a fruit juice or gasoline mixture that depend on the proportions of components in the product. They also form the starting point for Diversity-Interactions models used to study the relationships between ecosystem properties such as total biomass and community composition (Connolly et al., 2013). However, a CTI has a completely different nature to the kinds of properties typically modelled by Scheffé polynomials. Unlike, for example, the combustion properties of a gasoline mixture, a CTI cannot be directly measured, is not experienced by any organism in the community, and does not affect any property of the physical world. We are therefore free to choose the form of a CTI so that it has desirable properties. In particular, when used as a measure of change in a community, we would like a CTI to preserve the algebraic properties of community dynamics, so that we can understand what it means. A similar argument has been made that a community-weighted mean value of some property such as stomatal space use efficiency should be based on the geometric rather than the arithmetic mean, to reflect the multiplicative nature of organism growth (Liu et al., 2026).

Here, we will show that the algebraic properties of community dynamics that should be preserved by a CTI are those of the Aitchison geometry of compositional data (Pawlowsky-Glahn et al., 2015, chapter 3). The Aitchison geometry is the standard setting for statistical analysis of compositional data and is obviously relevant to relative abundance data (Billheimer et al., 2001), yet has been relatively little used by ecologists. The required algebraic properties determine a form of CTI different from those in common use. However, when relative abundances (or changes in relative abundances) are approximately equal and STIs are close to their geometric mean, we show that the commonest form of CTI approximately preserves the required algebraic properties. Working in the Aitchison geometry leads naturally to a classification of species as relative warm- or cold-affinity, in a way that does not depend on relative abundances, and to species contributions’ to a change in CTI over time that are consistent with the principles of population dynamics. Working in the Aitchison geometry also leads to a measurement of the extent to which change in relative abundances is in the direction given by the STIs, analogous to the *R*^2^ in a multiple regression. Calculating this quantity is an important check on whether a CTI contains useful information.

### Desirable properties for a CTI

Throughout, we will assume that a real-valued CTI is desired. To be a useful measure of change, a CTI should be:

**I1**. *Intensive, i*.*e. independent of the total size of the system;*

**I2**. *Additive with respect to consecutive changes;*

**I3**. *Homogeneous, i*.*e. proportional to the interval of time or space over which a constant rate of change occurs*.

These axioms have straightforward ecological motivations. It seems natural to want a CTI not to depend on the total size of the system (axiom I1), but on the relative abundances of each species, because multiplying the abundance of every species by the same constant does not indicate any change in the temperature preferences of individuals making up the community. The need for axioms I2 and I3 comes from the desire to interpret trends in the CTI over time or space. For example, Devictor et al. (2008) report and interpret a linear temporal trend in the CTI of French birds. Axiom I2 tells us that the same amount of change in a community leads to the same amount of change in the CTI, and axiom I3 tells us that if the rate of change in the community is constant over time, the amount of change in the index is proportional to the amount of time elapsed. Unless I2 and I3 are satisfied, we cannot interpret a linear trend in the CTI as a linear trend in the community.

The original CTI (Equation 1) clearly satisfies axiom I1, although some forms of CTI do not (supporting information, section S4). A function of abundances that is independent of the total size of the system (axiom I1) is sometimes called an intensive function (Tolman, 1917), and is homogeneous of degree zero with respect to abundances. In other words, it is constant along rays from the origin in the space 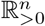 of abundances. These rays are the equivalence classes of compositions (Pawlowsky-Glahn et al., 2015, Definition 2.2): along any such ray, the relative abundances are constant. Therefore a CTI *I* must be a function *I*(**x**) of the *n* relative abundances alone. Relative abundances are compositional data, for which it is natural to work in the Aitchison geometry, a finite-dimensional inner product space over the reals, in which the vectors are elements of the standard (*n* − 1)-dimensional open simplex Δ^*n−*1^ (Pawlowsky-Glahn et al., 2015, chapter 3). Axioms I2 and I3 together mean that a CTI should be a homomorphism from the relative abundances to the real line, preserving the algebraic structure described below. That measurements should be homomorphisms is central to the relational theory of measurement (Wolff, 2020, chapter 5), although in theories of measurement there is additional structure imposed by an order relation on the objects to be measured, which we do not have for relative abundances.

### The Aitchison CTI

The Aitchison geometry (Pawlowsky-Glahn et al., 2015, section 3.2) has addition operation *perturbation*, denoted ⊕ and defined by

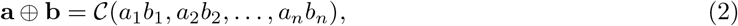

for **a, b** ∈ Δ^*n−*1^, where C(**a**) is the *closure* of the vector **a**:

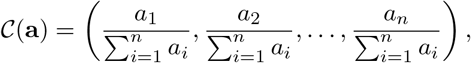

and scalar multiplication operation *powering*, denoted ⊙ and defined by

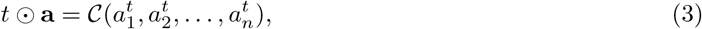

for *t* ∈ R and **a** ∈ Δ^*n−*1^.

The Aitchison geometry is the natural choice for studying the dynamics of relative abundances, because constant proportional changes correspond to constant perturbations, and the same proportional rate of change can be applied to different time intervals by powering (Pawlowsky-Glahn et al., 2015, section 9.1). In population dynamics, constant proportional change occurs when all aspects of the environment for each species are constant, analogous to constant velocity of a body with no forces acting on it (Ginzburg, 1986). Under constant proportional change, if the *i*th species has absolute abundances (measured in any appropriate dimensions) *y*_*i*_(0) and *y*_*i*_(1) at times 0 and 1 respectively, then

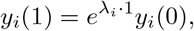

where 1 represents a unit of time (dimensions T) and *λ*_*i*_ is the proportional population growth rate of the *i*th species over this time interval (dimensions T^*−*1^). It follows immediately from the definition of perturbation (Equation 2) that the dynamics of relative abundances **x** = *C*(*y*_1_, *y*_2_, …, *y*_*n*_) are given by

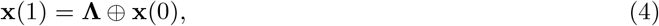

Where 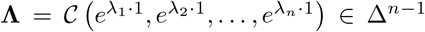. Note that changes in relative abundances can still be written in this form if the rates of change are not constant: the *λ*_*i*_ may instead represent mean rates over the time interval. Over consecutive time intervals 1, 2 with corresponding vectors of change **Λ**_1_, **Λ**_2_, repeated application of Equation 4 gives the total amount of proportional change **Λ**_2_ ⊕ **Λ**_1_. Then axiom I2 means that we require

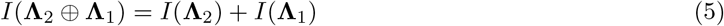

for all **Λ**_2_, **Λ**_1_ ∈ Δ^*n−*1^. Applying a constant proportional rate over some time interval *t*, it follows immediately from the definition of powering (Equation 3) that relative abundances **x**(*t*) at time *t* are given by

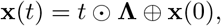

where *t* is written as a pure number, because one unit of time appears in the rates **Λ**. Then axiom I3 means that we require

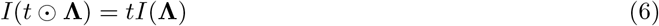

for all *t* ∈ R and all **Λ** ∈ Δ^*n−*1^.

Together, I1, I2 and I3 mean that a CTI must be a linear functional from Δ^*n−*1^ to R (Axler, 2015, Definition 6.39). By the Riesz representation theorem (Axler, 2015, Theorem 6.42), for any such linear functional there is a unique vector **c** ∈ Δ^*n−*1^ such that for all **x** ∈ Δ^*n−*1^, *I*(**x**) = ⟨**x, c**⟩ _*a*_, where ⟨ ·,· ⟩_*a*_ denotes the Aitchison inner product (Pawlowsky-Glahn et al., 2015, section 3.3). We therefore make the following definition:

#### Definition 1 (Aitchison CTI)

*The Aitchison CTI with parameter* **c**, *I*_**c**_(**x**), *is the Aitchison inner product of the relative abundances* **x** *with a fixed vector* **c** *of STIs*,

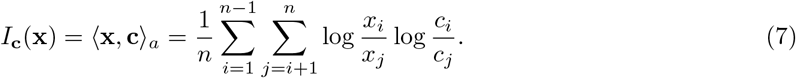

From the properties of an inner product (Axler, 2015, Definition 6.3), we know that any choice of fixed **c** in Equation 7 will give us a linear functional. The obvious choice is to take **c** to be the STIs (measured in an absolute temperature scale such as Kelvin so that the ratios *c*_*i*_*/c*_*j*_ are meaningful). Because the STIs *c*_*i*_ appear only as ratios, the Aitchison CTI is dimensionless and is affected only by relative STIs. Also note that since a vector of change in relative abundances **Λ** is an element of Δ^*n−*1^, the corresponding change in the Aitchison CTI is given by *I*_**c**_(**Λ**) The behaviour of the Aitchison CTI is easy to understand. In the following explanation, we break down the ways in which a pair of species (*i, j*) contribute to the Aitchison CTI. We refer to relative abundances, but throughout, the same explanations apply to changes in relative abundance. We also assume that the STIs *c*_*i*_ are not all equal (if they were, then every set of relative abundances would give a zero CTI). If the relative abundance *x*_*i*_ of species *i* is greater than the relative abundance *x*_*j*_ of species *j*, and the STI *c*_*i*_ for *i* is greater than the STI *c*_*j*_ for *j*, then both log(*x*_*i*_*/x*_*j*_) and log(*c*_*i*_*/c*_*j*_) will be positive, and the pair (*i, j*) will make a positive contribution to the Aitchison CTI. Similarly, if *x*_*i*_ *< x*_*j*_ and *c*_*i*_ *< c*_*j*_ (species *i* has both lower relative abundance and lower STI than species *j*), then the pair (*i, j*) will make a positive contribution. Conversely, if *x*_*i*_ *> x*_*j*_ and *c*_*i*_ *< c*_*j*_, or *x*_*i*_ *< x*_*j*_ and *c*_*i*_ *> c*_*j*_, then the pair (*i, j*) will make a negative contribution. The pair (*i, j*) makes a zero contribution if and only if either *x*_*i*_ = *x*_*j*_ (the species have the same relative abundance) or *c*_*i*_ = *c*_*j*_ (the species have the same STI). However, there are many ways to get *I*_**c**_ = 0, even if no pair makes a zero contribution. The null space, null *I*_**c**_, of the Aitchison CTI is the set of relative abundances for which the Aitchison CTI is equal to zero. Since dim Δ^*n−*1^ = *n* − 1 but dim ℝ = 1, *I*_**c**_ cannot be injective if we have more than two species (Axler, 2015, Theorem 3.23), and therefore null *I*_**c**_ does not consist only of the zero vector **0** = (1*/n*, …, 1*/n*) in the Aitchison geometry (Axler, 2015, Theorem 3.16). By the Fundamental Theorem of Linear Maps (Axler, 2015, Theorem 3.22),

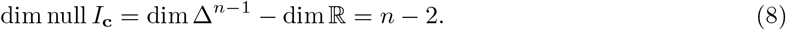

Thus the subspace of relative abundances with zero Aitchison CTI has dimension *n*− 2. This is particularly important when looking at change in the Aitchison CTI. Equation 8 tells us that there is a sense in which “most” changes in relative abundance do not change the Aitchison CTI.

### Relative warm- and cold-affinity species, and contributions to change

Warm- and cold-affinity species have been defined as species whose STI is higher or lower than the DeVictor CTI for the community respectively (McLean et al., 2021), so that the Devictor CTI will increase if the relative abundance of a warm-affinity species increases and decrease if the relative abundance of a cold-affinity species increases. However, McLean et al. (2021) note that whether a species is warm- or cold-affinity depends on the relative abundances of all species in the community, not just on the STIs. This is obviously undesirable, because it requires an arbitrary choice of a reference CTI, such as the mean CTI for a location over a given time period (McLean et al., 2021) or the CTI for a location at the start of a time period (Dobson et al., 2026).

We would like a definition of warm- and cold-affinity species that does not depend on relative abundances. Such a definition arises naturally for the Aitchison CTI. We use the terms relative warm- and cold-affinity, because both relative abundances and STIs appear only as ratios in Equation 7. The Aitchison CTI (Equation 7) can be rewritten in the form

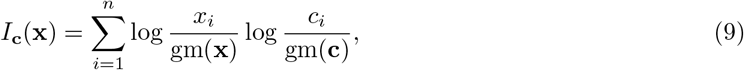

where gm(·) denotes the geometric mean (Pawlowsky-Glahn et al., 2015, section 3.3). In this form, it appears that if a species has an STI greater than the geometric mean of the STIs for all species in the community, then an increase in its relative abundance will be associated with an increase in the Aitchison CTI, and vice versa. However, some care is needed because we cannot increase the relative abundance of the *i*th species without changing the relative abundances of other species in some way. Thus we need an appropriate definition of the partial derivative for a composition. Let the part C-derivative of *I*_**c**_ with respect to *x*_i_, denoted 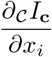, be the rate of change of *I*_c_ under a multiplicative increase in *x*, while the ratios of all other relative abundances are held constant (Barceló-Vidal et al., 2011). Such a pattern of change is the natural choice because it leaves the subcomposition consisting of relative abundances of all species other than *i* unchanged. We show in the supporting information, section S5, that this part C-derivative is given by

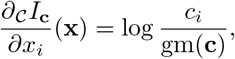

which does not depend on **x**, is positive if species *i* has an STI greater than the geometric mean of the STIs for all species in the community, and is negative if species *i* has an STI less than the geometric mean of the STIs for all species in the community. Thus we define a relative warm-affinity species as a species whose STI is greater than the geometric mean of STIs for all species in the community, and a relative cold-affinity species as a species whose STI is less than the geometric mean of the STIs for all species in the community.

McLean et al. (2021) proposed partitioning contributions to the Devictor CTI into tropicalization (increases in warm-affinity species), deborealization (decreases in cold-affinity species), borealization (increases in cold-affinity species) and detropicalization (decreases in warm-affinity species). However, their partitioning is not exact because their definitions of warm- and cold-affinity species depend on relative abundances. Similarly, Dobson et al. (2026) plotted differences in relative abundance between warmed and ambient treatments against differences between species’ STIs and the CTI at an arbitrary time point. In both cases, the use of a CTI that does not satisfy axioms I2 and I3 makes it difficult to interpret the results. However, we can adapt the Dobson et al. (2026) approach to the Aitchison geometry to visualize the contributions that proportional changes in the relative abundance of relative warm- and cold-affinity species make to the Aitchison CTI. It is straightforward to show from Equation 9 that the change in Aitchison CTI between two time points is given by

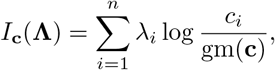

where *λ*_*i*_ = log(*x*_*i,t*_*/x*_*i*,0_) is the proportional change in the relative abundance of the *i*th species between times 0 and *t* (treating *t* as a unit of time). Thus the *i*th species makes contribution *λ*_*i*_ log(*c*_*i*_*/*gm(**c**)) to the change in Aitchison CTI, and we can plot *λ*_*i*_ against log(*c*_*i*_*/*gm(**c**)) for each species, overlaying contours of the contribution *λ*_*i*_ log(*c*_*i*_*/*gm(**c**)). Those species in the top right and bottom left corners of the plot make positive contributions to the change in Aitchison CTI, and those species in the top left and bottom right corners make negative contributions. Note that these contributions cannot be independent, because there are *n* species but Δ^*n−*1^ is (*n*− 1)-dimensional. Nevertheless, such a plot may help us to understand what causes a change in Aitchison CTI. In particular, a doubling of relative abundance will make the same contribution to change in the Aitchison CTI, whether the species is rare or common, for a given relative STI. This is consistent with the principles of population dynamics discussed above.

### Example: Devictor and Aitchison CTIs under exponential growth

We illustrate the behaviour of the standard Devictor CTI (Equation 1) and the Aitchison CTI (Equation 7) with a simple example. Consider a set of three species with initial abundances **y**(0) = (3, 2, 1) in some units of abundance, growing exponentially with proportional population growth rates ***λ*** = (− 0.2, 0.1, 0.25) per some unit of time, and STIs **c** = (283.15 K, 293.15 K, 288.15 K). Because each species is growing exponentially, log abundances are linear functions of time (Figure 1a). The relative abundances are also linear functions of time in the Aitchison geometry (Equation 4), but not in ℝ ^3^ with the usual vector addition and scalar multiplication operations (Figure 1b). Thus the standard CTI is not a linear function of time in this case (Figure 1c). An ecologist would be tempted to incorrectly conclude that something has changed around time *t* = 7, where the CTI changes from increasing to decreasing. In contrast, the Aitchison CTI increases over time at a constant rate (Figure 1d), correctly indicating that there has been no change in the dynamics of the community.

**Figure 1:**
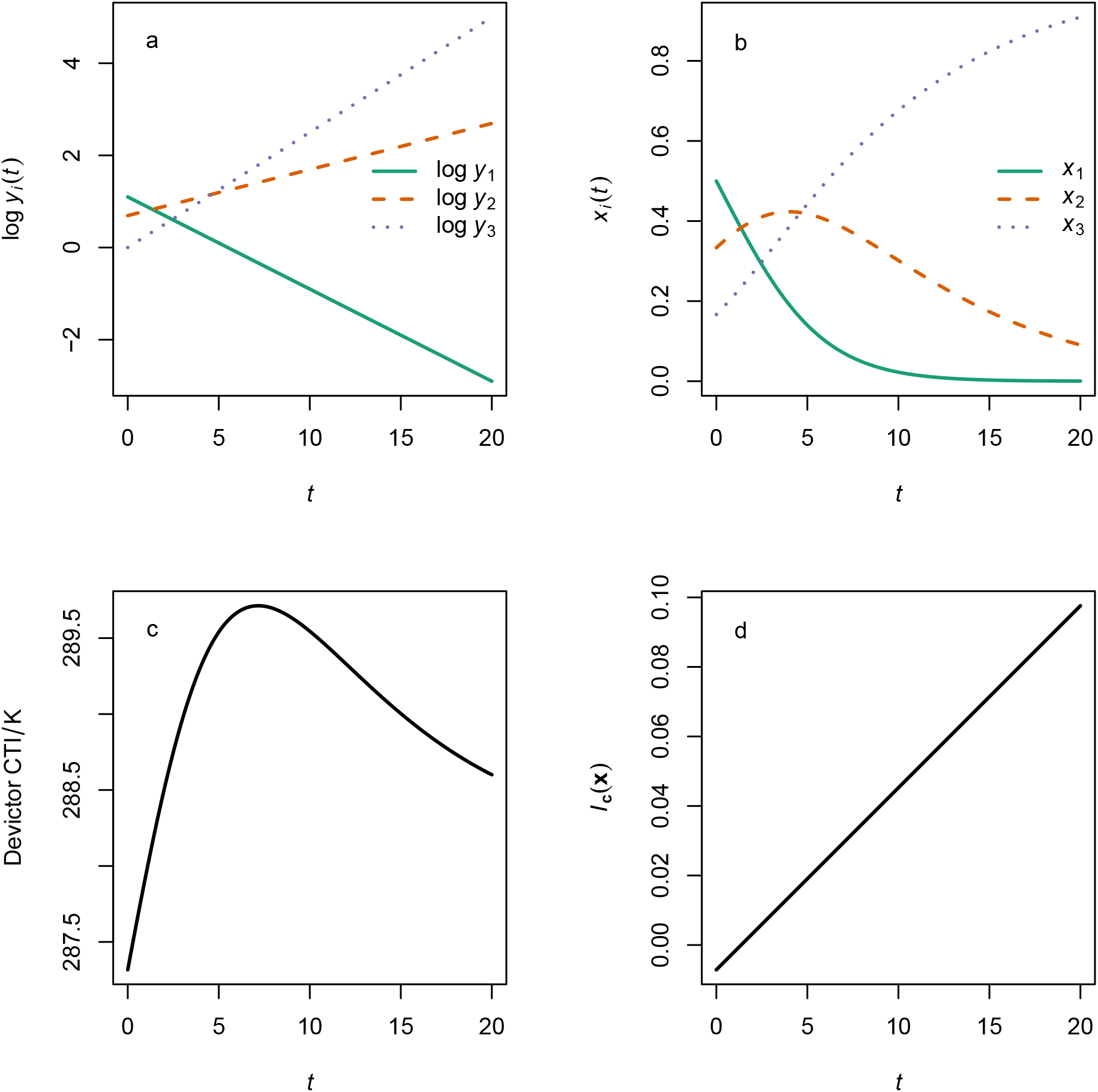
Behaviour of the standard Devictor Community Temperature Index (Equation 1) and the Aitchison Community Temperature Index (Equation 7) in a simple case: (a) log abundances against time for three species, each with constant proportional population growth rate; (b) relative abundances against time for these three species; (c) the standard Devictor Community Temperature Index (with units K) against time; (d) the Aitchison Community Temperature Index (dimensionless) against time. Initial abundances **y**(0) = (3, 2, 1) in some units of abundance, proportional population growth rates ***λ*** = (−0.2, 0.1, 0.25) per some unit of time, Species Temperature Indices **c** = (283.15 K, 293.15 K, 288.15 K).

In this example, the geometric mean STI is 288.12 K, so species 1 is a relative cold-affinity species, while species 2 and 3 are relative warm-affinity species. As noted above, this classification does not depend on the relative abundances.

### Approximate relationship between the Aitchison CTI and the original CTI

In some cases, there may be a good reason for using the original Devictor CTI (Equation 1). Since this original CTI is not an Aitchison inner product, it cannot satisfy axioms I2 and I3, making interpretation of patterns of change difficult. However, we show in the supporting information, section S6, that provided relative abundances (or changes in relative abundances) are approximately equal, and STIs are close to their geometric mean, the relationship between the Aitchison CTI and the original CTI is approximately linear with slope *n/*gm(**c**) and intercept −*n*. Thus the original CTI may approximately have the properties we want from a measure of change in a community.

### Extent to which change in relative abundances is in the direction of the relative STIs

We would like to be able to measure the extent to which change in relative abundances is in the direction of the relative STIs, or in other words, how much the CTI is telling us about community dynamics. Working with the Aitchison CTI allows us to do this through a straightforward generalization of the coefficient of determination, *R*^2^. In an ordinary multiple regression in ℝ^*n*^, *R*^2^ can be defined geometrically as the square of the cosine of the angle between the response vector and the subspace spanned by the explanatory vectors (Saville and Wood, 1991, p. 446). This definition generalizes to an arbitrary finite-dimensional inner product space. In particular, in the Aitchison geometry, we can define the square of the cosine of the angle *θ* between the vector **c** and the space spanned by a set of *k* vectors of change in relative abundance, span(**Λ**_1_, **Λ**_2_, …, **Λ**_*k*_), as

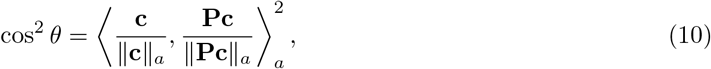

(Eaton, 2007, pp. 61-62), where ∥ · ∥_*a*_ denotes the norm induced by the Aitchison inner product (Pawlowsky-Glahn et al., 2015, section 3.3), and **P** is the orthogonal projection onto span(**Λ**_1_, **Λ**_2_, …, **Λ**_*k*_) given by **P** = **Z**(**Z**^T^**Z**)^*−*1^**Z**^T^, where **Z** is a matrix with columns **Λ**_1_, **Λ**_2_, …, **Λ**_*k*_ (Eaton, 2007, chapter 4). The necessary matrix operations in the Aitchison geometry can be defined in terms of isometric logratio transformation to ℝ^*n−*1^, standard matrix operations in ℝ^*n−*1^ and back-transformation (Pawlowsky-Glahn et al., 2015, sections 4.4, 4.9), but in practice it is easier to transform to ℝ^*n−*1^ and do the computations there, since we only need an answer in ℝ. Thus Equation 10 gives a measure of the extent to which change in relative abundances is in the direction of the relative STIs, taking values between 0 and 1, with 0 if the vector of relative STIs is orthogonal to the subspace spanned by the vectors of change in relative abundances, and 1 if the vector if relative STIs is in this subspace.

### Example: macrobenthos in the Bay of Biscay

We illustrate the relationship between the Aitchison CTI and the original CTI, the extent to which change in relative abundances is in the direction of the relative STIs, and contributions from each species, using data on the hard-substrate macrobenthos community of the Bay of Biscay between 2002 to 2020 (Chust et al., 2024a, supporting information for data set 12). These data are provided in Chust et al. (2024b). The data contain estimated abundances of 151 species of macroinvertebrates, lichens and macroalgae, obtained by averaging values from a semi-quantitative 7-point scale over multiple transects. Over all species and years, 51% of estimated abundances were zero. We follow Chust et al. (2024a) in dealing with zeros by adding 1 to all observed abundances, although the largest observed value was only 5. The data set also contains STIs, obtained by matching occurrence records with sea surface temperature records for each species (Chust et al., 2024a). Computation is straightforward using the R package compositions (van den Boogaart et al., 2025) and R version 4.5.2 (R Core Team, 2025). Code and data are available at Spencer (2026).

In these data, the Aitchison CTI is approximately linearly related to the original CTI (Figure 2), and it may therefore be reasonable to interpret patterns of change in the original CTI over time. Some caution is needed given the large proportion of zeros, meaning that the estimated relative abundances may be artificially close to the zero vector in the Aitchison geometry about which the approximation is made (supporting information, section S6). The requirement that all the STIs are fairly close to their geometric mean appears to be met (range 283 K to 297K, geometric mean 288 K), unsurprisingly for a set of species living in the same location. However, the value of cos^2^ *θ* is only 0.12, which suggests that the CTI is giving us only a modest amount of information about community dynamics. It would be surprising if the dynamics of a community of 151 species could be explained adequately in one dimension. Changes in relative abundance in this community live in the 150-dimensional simplex, but Equation 8 tells us that there is a 149-dimensional subspace of changes in relative abundance that do not change the Aitchison CTI. Furthermore, the large proportion of zeros may mean that the abundances of many species at many time points are poorly estimated, which may reduce the amount of information about community dynamics that can be obtained.

**Figure 2:**
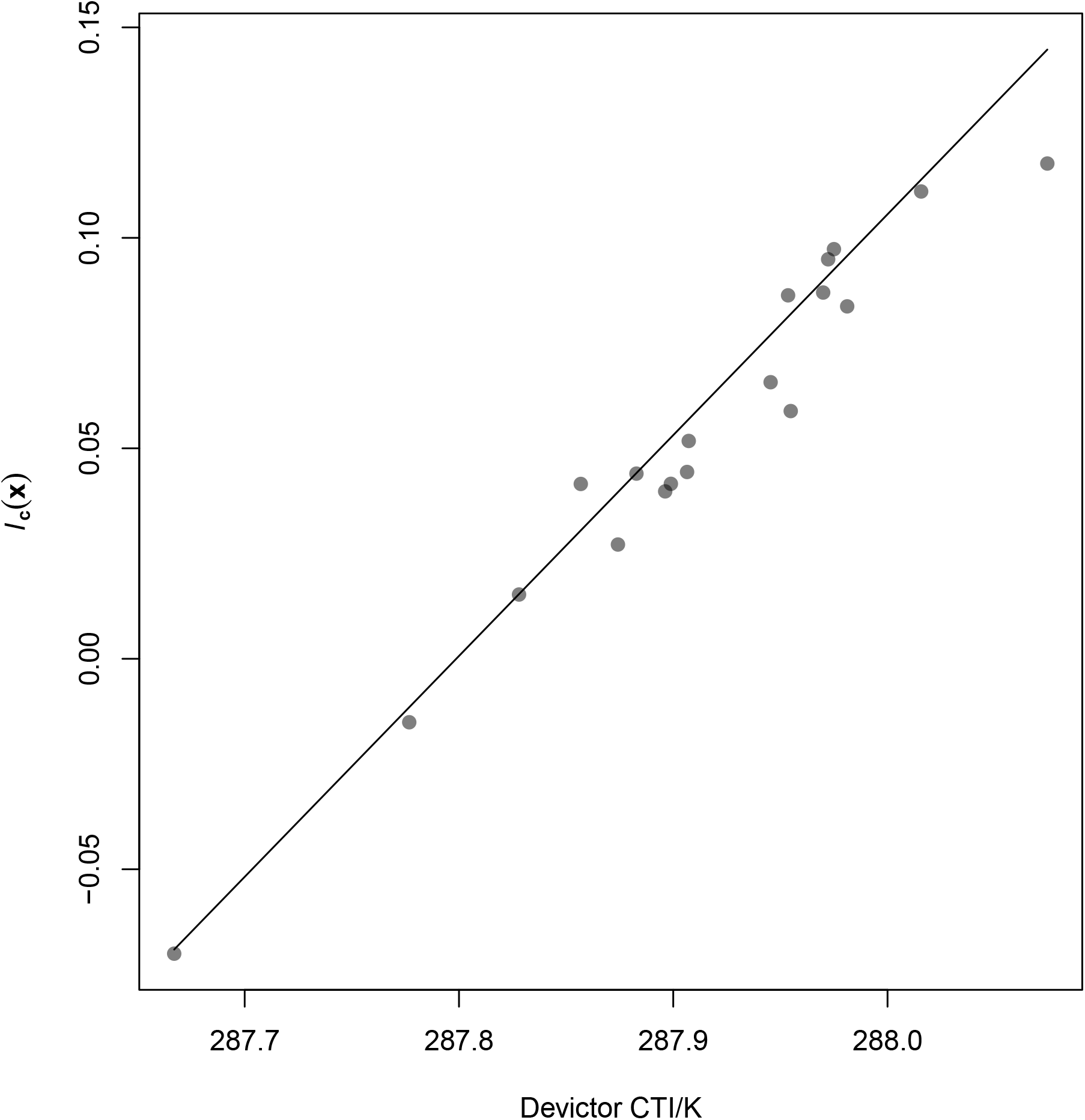
Relationship between the Aitchison CTI *I*_**c**_(**x**) (Equation 7, dimensionless) and the original Devictor CTI (Equation 1, in Kelvin), for data from the hard-substrate macrobenthos community in the Bay of Biscay between 2002 and 2020 (Chust et al., 2024a), with 1 added to all abundances before analysis to deal with zeros. Each point represents data from a single year. The line is the linear function of the Devictor CTI with slope *n/*gm(**c**) and intercept − *n* that approximates the Aitchison CTI (supporting information, section S6).

In the Bay of Biscay data, most species made small contributions to the change in Aitchison CTI between 2002 and 2020 (Figure 3: the horizontal line of points with *λ*_*i*_ just below zero corresponds to species with zero observed values in both 2002 and 2020, since the mean abundance increased between 2002 and 2020). There was no obvious systematic relationship between relative increases or decreases in abundance and relative temperature preferences. The extreme outlying contributions from a few species to change in CTI in response to experimental warming found by Dobson et al. (2026) in North American plant communities were not observed in these data. However, the results are not directly comparable, because Dobson et al. (2026) did not use the Aitchison CTI, their data were experimental rather than observational, and the Bay of Biscay data may have poorly-estimated relative abundances for many species.

**Figure 3:**
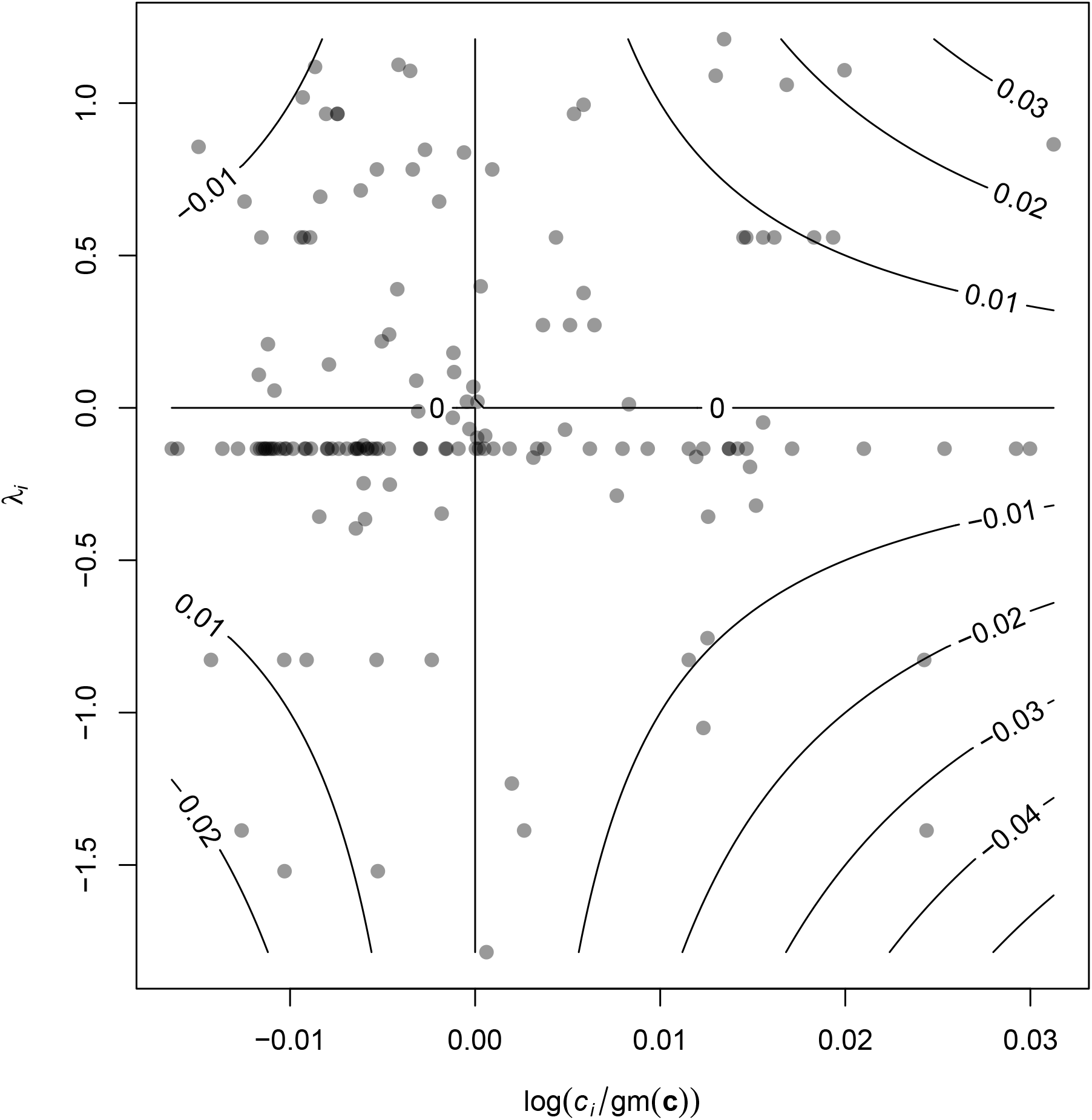
Species contributions to the change in Aitchison CTI *I*_**c**_(**x**) (Equation 7, dimensionless) between 2002 and 2020, for data from the hard-substrate macrobenthos community in the Bay of Biscay (Chust et al., 2024a), with 1 added to all abundances before analysis to deal with zeros. Each point represents the proportional change in relative abundance *λ*_*i*_ = log(*x*_*i*,2020_*/x*_*i*,2002_) and the relative STI log(*c*_*i*_*/*gm(**c**)) for a single species. The product *λ*_*i*_ log(*c*_*i*_*/*gm(**c**)) (contours) is the contribution that the species makes to the change in Aitchison CTI.

### Recommendations

We end with four recommendations for users of community temperature indices.

1. Report which form of CTI was used using mathematical notation rather than words alone, to avoid ambiguity.
2. If studying changes in a CTI over time or space, use the Aitchison CTI (with temperatures in an absolute scale such as the Kelvin scale) unless there is a compelling reason not to, because the Aitchison CTI is the only form that preserves the algebraic properties of community dynamics. The Aitchison CTI also leads naturally to a definition of relative warm- and cold-affinity species that does not depend on relative abundances, and to contributions to change in CTI that are consistent with the principles of population dynamics.
3. If there is a compelling reason to use some other form of CTI to study changes over time or space, check whether there is an approximately linear relationship between this form and the Aitchison CTI, so that patterns in the chosen CTI are at least approximately interpretable.
4. If using the Aitchison CTI, calculate the extent to which change in relative abundances is in the direction of the relative STIs. Doing so will give some indication of whether the CTI is describing a meaningful pattern of change in relative abundances.

## Acknowledgements

I am grateful to Guillem Chust and Ernesto Villarino for explaining details of the Bay of Biscay data, and to Guillem Chust for comments on the manuscript.

