## Supporting information for "The algebra of community temperature indices"

### The algebra of community temperature indices: supporting information

Matthew Spencer

August 2, 2026

#### S1 Literature search terms and methods

I searched Web of Science (`https://www.webofscience.com`) using the term "community temperature index" on 26 August 2025. Clearly this search will not retrieve all papers calculating a community temperature index (CTI), but is likely to be sufficient to give a rough idea of the approaches used. 93 papers were found (1). I read each paper to determine what form of CTI was calculated, as described below (in some cases, more than one form was used within a single paper). Papers for which no CTI was calculated were excluded from further analysis, but are listed in Table 1 with the forms used column blank. For each paper, I also recorded whether changes in CTI over time and/or space were studied. I included differences between locations, habitat types and effects of environmental variables associated with locations, such as elevation and protection status, in changes over space. Search results and code are available at Spencer (2026).

#### S2 Forms of community temperature index

I determined the form of CTI used from the description in the methods section of each paper, although supporting information was also checked if explicitly referred to and the methods section was ambiguous. In many cases, the form of CTI was described in words rather than mathematical notation, and in some cases the words were unclear or contradictory. In such cases, I have given my best guess at what was intended. Papers that stated they calculated an abundance-weighted CTI were assumed to use form 1 below unless a more explicit statement was given that contradicted this. Papers that stated they calculated the mean of Species Temperature Indices (STIs) across species were assumed to use form 3 below unless a more explicit statement was given that contradicted this. I took "weighted by" to mean a set of weights (such as relative abundances) that sum to 1, unless a formula was explicitly given that stated otherwise.

In the equations below, $n$ denotes the number of species, $c_i$ denotes the STI for the $i$th species, $x_i$ denotes the relative abundance of the $i$th species, $y_i$ denotes the absolute abundance of the $i$th species, $\mathbf{1}_{y_i>0}$ denotes the indicator function taking the value 1 if the $i$th species is present and 0 if it is absent, and $p_i$ is the occupancy of the $i$th species (the proportion of locations at which it is present). I did not distinguish between different forms of abundance such as counts, model-based estimates and ordered categories, and did not distinguish abundance from related quantities such as biomass and basal area, because these distinctions do not affect the form of the CTI, although they may affect its interpretation.

The following forms of CTI appear to have been used.

$$\sum_{i=1}^{n} c_i x_i = \frac{\sum_{i=1}^{n} c_i y_i}{\sum_{i=1}^{n} y_i} \tag{1}$$

$$\frac{\sum_{i=1}^{n} c_i \log y_i}{\sum_{i=1}^{n} \log y_i} \quad \text{or} \quad \frac{\sum_{i=1}^{n} c_i \log(y_i+1)}{\sum_{i=1}^{n} \log(y_i+1)} \tag{2}$$

$$\frac{\sum_{i=1}^{n} c_i \mathbf{1}_{y_i>0}}{\sum_{i=1}^{n} \mathbf{1}_{y_i>0}} \tag{3}$$

$$\frac{\sum_{i=1}^{n} c_i p_i}{\sum_{i=1}^{n} p_i} \tag{4}$$

Note that the equations given in Villarino et al. (2020) and Chust et al. (2024) are shorthand notations for form 2 (E. Villarino, pers. comm).

#### S3 Literature search results

Of the 93 papers found (Table 1), 2 were excluded because no CTI was calculated. Of the remaining 91 papers, 53 (58%) used form 1, 11 (12%) used form 2, 36 (40%) used form 3 and 2 (2%) used form 4. Of the 91 papers, 80 (88%) studied some aspect of change in CTI over time, and 84 (92%) studied some aspect of change in CTI over space.

Table 1: Literature search for "community temperature index" on Web of Science, as described in Section S1. Numbers in the forms used column correspond to equation numbers in Section S2, as far as could be determined from descriptions in the papers. The forms used column is blank if no CTI was calculated in the paper. In the change over time and change over space columns, a y indicates that some aspect of change in CTI over time or space respectively was studied. Change over space includes differences between locations and effects of associated environmental variables.

| | source | forms used | change over time | change over space |
|---|---|---|---|---|
| 1 | Devictor et al. (2008) | 1 | y | y |
| 2 | Godet et al. (2011) | 2, 3 | y | |
| 3 | Kampichler et al. (2012) | 1 | y | y |
| 4 | Filz et al. (2013) | 3 | y | y |
| 5 | Lindström et al. (2013) | 1, 3 | y | y |
| 6 | Reif et al. (2013) | 1 | y | |
| 7 | Schweiger et al. (2014) | | | |
| 8 | Zografou et al. (2014) | 1 | y | y |
| 9 | Princé and Zuckerberg (2015) | 1, 3 | y | y |
| 10 | Savage and Vellend (2015) | 1 | y | y |
| 11 | Pearce-Higgins et al. (2015) | | | |
| 12 | Nieto-Sánchez et al. (2015) | 1, 3 | y | y |
| 13 | Thomsen et al. (2016) | 3 | y | |
| 14 | Tayleur et al. (2016) | 1, 3 | y | y |
| 15 | Li et al. (2016) | 3 | y | y |
| 16 | Martay et al. (2016) | 1 | y | y |
| 17 | Stuart-Smith et al. (2017) | 2 | y | y |
| 18 | Santangeli and Lehikoinen (2017) | 1 | y | y |
| 19 | Gaüzère et al. (2017) | 1 | y | y |
| 20 | Santangeli et al. (2017) | 1 | y | y |
| 21 | Oliver et al. (2017) | 1 | y | y |
| 22 | Bates et al. (2017) | 3 | y | y |
| 23 | Bowler and Böhning-Gaese (2017) | 1 | y | |
| 24 | Kwon (2017) | 1, 3 | | y |
| 25 | Sparrius et al. (2018) | 1 | | y |
| 26 | Gaget et al. (2018) | 2 | y | y |
| 27 | Flanagan et al. (2019) | 3 | y | y |
| 28 | Fourcade et al. (2019) | 3 | y | y |
| 29 | Haase et al. (2019) | 1, 3 | y | y |
| 30 | Cerrato et al. (2019) | 3 | y | y |
| 31 | Termaat et al. (2019) | 4 | y | y |
| 32 | Brice et al. (2019) | 1 | y | y |
| 33 | Löffler et al. (2019) | 3 | y | y |
| 34 | Burrows et al. (2020) | 1 | y | y |
| 35 | Fumy et al. (2020) | 3 | y | y |
| 36 | Villarino et al. (2020) | 2 | y | y |

Table 1: continued

| | source | forms used | change over time | change over space |
|---|---|---|---|---|
| 37 | Gaget et al. (2020) | 2 | y | y |
| 38 | Assandri (2021) | 3 | y | y |
| 39 | Dietz et al. (2020) | 3 | y | y |
| 40 | Haubrock et al. (2020) | 1 | y | |
| 41 | Ajani et al. (2020) | 1 | y | |
| 42 | Gaget et al. (2021a) | 1 | y | y |
| 43 | Popović et al. (2021) | 1 | | y |
| 44 | Lehikoinen et al. (2021b) | 1 | y | y |
| 45 | Gaget et al. (2021b) | 3 | y | y |
| 46 | Bonachela et al. (2021) | 1 | | y |
| 47 | Lehikoinen et al. (2021a) | 1 | y | y |
| 48 | Fourcade et al. (2021) | 1, 3 | y | y |
| 49 | Kwon et al. (2021) | 4 | y | y |
| 50 | Mingarro et al. (2021) | 3 | y | y |
| 51 | Richard et al. (2021) | 3 | y | y |
| 52 | Freeman et al. (2021) | 1 | y | y |
| 53 | Santorufo et al. (2021) | 1 | y | y |
| 54 | Comte et al. (2021) | 2 | y | y |
| 55 | Fartmann et al. (2021) | 3 | y | y |
| 56 | McLean et al. (2021) | 1 | y | y |
| 57 | Christiansen et al. (2022) | 3 | y | y |
| 58 | Batten et al. (2022) | 1 | y | y |
| 59 | Scott et al. (2022) | 1 | y | y |
| 60 | Schuster et al. (2022) | 3 | | y |
| 61 | Curley et al. (2022) | 1 | y | y |
| 62 | Fartmann et al. (2022) | 3 | y | y |
| 63 | Lajeunesse and Fourcade (2023) | 3 | y | y |
| 64 | Soler et al. (2022) | 2 | y | y |
| 65 | de Souza and dos Santos (2023) | 1 | y | y |
| 66 | Borderieux et al. (2023) | 3 | y | y |
| 67 | Gaget et al. (2024) | 2 | y | y |
| 68 | Dietrich et al. (2023) | 1 | y | |
| 69 | Arriaga et al. (2023) | 1 | y | y |
| 70 | de Azevedo et al. (2023) | 1 | y | y |
| 71 | Hintsanen et al. (2023) | 1 | y | y |
| 72 | Marjakangas et al. (2023) | 1 | y | y |
| 73 | Kwon et al. (2024) | 3 | y | y |
| 74 | Álvarez et al. (2024) | 3 | y | y |
| 75 | De Pauw et al. (2024) | 1 | | y |
| 76 | Finegan et al. (2024) | 1 | | y |
| 77 | Arriaga et al. (2024b) | 1 | y | y |
| 78 | Gril et al. (2024) | 1, 3 | | y |
| 79 | Chust et al. (2024) | 2 | y | y |
| 80 | Goßmann et al. (2024) | 3 | | y |
| 81 | Pacheco-Riaño et al. (2024) | 3 | y | y |
| 82 | Aston et al. (2024) | 2 | | y |
| 83 | Arriaga et al. (2024a) | 1 | y | y |
| 84 | Hemberger and Williams (2024) | 1, 3 | y | y |
| 85 | Pessarrodona et al. (2024) | 1 | y | y |
| 86 | Ursul et al. (2025) | 3 | y | y |
| 87 | Hintsanen et al. (2025) | 1 | y | y |
| 88 | Chen et al. (2025) | 1 | y | y |
| 89 | Mäkinen et al. (2025) | 1 | y | y |
| 90 | Verniest et al. (2025) | 1, 3 | y | y |
| 91 | Gossmann et al. (2025) | 1 | | y |

Table 1: continued

| | source | forms used | change over time | change over space |
|---|---|---|---|---|
| 92 | Jonas et al. (2025) | 2 | y | y |
| 93 | Alba and Chamberlain (2025) | 1 | y | y |

#### S4 Not every CTI that has been used is intensive

Here, we show that one of the forms that has appeared in the literature (Equation 2) is not intensive. We concentrate on the version using $\log(y_i)$ rather than $\log(y_i + 1)$ for simplicity. Suppose that all abundances $y_i$ are multiplied by the same positive constant $k$. Then this form, which we refer to as $\mathrm{CTI}_{\log}(\mathbf{y})$, where $\mathbf{y} = (y_1, y_2, \ldots, y_n)$, becomes

$$\begin{aligned}\mathrm{CTI}_{\log}(\mathbf{y}) &= \frac{\sum_{i=1}^{n} c_i \log(ky_i)}{\sum_{i=1}^{n} \log(ky_i)} \\ &= \frac{\log k \sum_{i=1}^{n} c_i + \sum_{i=1}^{n} c_i \log y_i}{n \log k + \sum_{i=1}^{n} \log y_i}.\end{aligned}$$

Differentiating with respect to $k$,

$$\frac{\mathrm{d}\mathrm{CTI}_{\log}(\mathbf{y})}{\mathrm{d}k} = \frac{\frac{1}{k} \sum_{i=1}^{n} c_i \sum_{i=1}^{n} \log y_i - \frac{n}{k} \sum_{i=1}^{n} c_i \log y_i}{(n \log k + \sum_{i=1}^{n} \log y_i)^2},$$

which will not be zero in general. Thus, multiplying all abundances by the same positive constant will change the value of form 2 of the CTI.

### S5 Part $\mathcal{C}$-derivatives of the Aitchison CTI

Let the part $\mathcal{C}$-derivative of $I_{\mathbf{c}}$ with respect to $x_i$, denoted $\frac{\partial_{\mathcal{C}} I_{\mathbf{c}}}{\partial x_i}$, be the rate of change of $I_{\mathbf{c}}$ under a multiplicative increase in $x_i$, while the ratios of all other relative abundances are held constant (Barceló-Vidal et al., 2011). Here, we calculate these part $\mathcal{C}$-derivatives.

First, note that we can rewrite the Aitchison CTI $I_{\mathbf{c}}(\mathbf{x})$ in the form:

$$I_{\mathbf{c}}(\mathbf{x}) = \langle \mathbf{x}, \mathbf{c} \rangle_a = \sum_{i=1}^{n} \log \frac{x_i}{\mathrm{gm}(\mathbf{x})} \log \frac{c_i}{\mathrm{gm}(\mathbf{c})},$$

where $\mathrm{gm}(\cdot)$ denotes the geometric mean (Pawlowsky-Glahn et al., 2015, section 3.3). By Proposition 13.3.5 in Barceló-Vidal et al. (2011),

$$\frac{\partial_{\mathcal{C}} I_{\mathbf{c}}}{\partial x_i}(\mathbf{x}) = x_i \left( \frac{\partial I_{\mathbf{c}}}{\partial x_i}(\mathbf{x}) - \sum_{j=1}^{n} x_j \frac{\partial I_{\mathbf{c}}}{\partial x_j}(\mathbf{x}) \right),$$

where $\frac{\partial I_{\mathbf{c}}}{\partial x_i}$ denotes the ordinary partial derivative, considering $I_{\mathbf{c}}(\mathbf{x})$ as a function in $\mathbb{R}^n$. We have

$$\frac{\partial I_{\mathbf{c}}}{\partial x_i} = \frac{1}{x_i} \log \frac{c_i}{\mathrm{gm}(\mathbf{c})},$$

and so

$$\begin{aligned} \frac{\partial_{\mathcal{C}} I_{\mathbf{c}}}{\partial x_i}(\mathbf{x}) &= x_i \left( \frac{1}{x_i} \log \frac{c_i}{\mathrm{gm}(\mathbf{c})} - \sum_{j=1}^{n} x_j \frac{1}{x_j} \log \frac{c_j}{\mathrm{gm}(\mathbf{c})} \right) \\ &= \log \frac{c_i}{\mathrm{gm}(\mathbf{c})}. \end{aligned} \tag{5}$$

#### S6 Approximately linear relationship between the Aitchison CTI and the original Devictor CTI

Here, we show that the Aitchison CTI is approximately linearly related to the original CTI from Devictor et al. (2008), which is also the most commonly-used CTI (1), if relative abundances are approximately equal (i.e. $nx_i \approx 1$ for $i = 1, \ldots, n$) and STIs are close to the geometric mean of STIs (i.e $c_i/\mathrm{gm}(\mathbf{c}) \approx 1$ for $i = 1, \ldots, n$).

Let $\mathbf{0} = (1/n, 1/n, \ldots, 1/n)$ denote the composition in which each species has equal relative abundance (the additive identity in the Aitchison geometry). Since the Aitchison CTI is a linear functional, it is by definition linear in $\mathbf{x}$ (Axler, 2015, Definition 6.39), and so using Definition 13.3.6 in Barceló-Vidal et al. (2011),

$$I_{\mathbf{c}}(\mathbf{x}) = I_{\mathbf{c}}(\mathbf{0}) + \mathbf{D}_{\mathcal{C}} I_{\mathbf{c}}(\mathbf{0}) \log \mathbf{x},$$

where

$$\mathbf{D}_{\mathcal{C}} I_{\mathbf{c}}(\mathbf{0}) = \left( \frac{\partial_{\mathcal{C}} I_{\mathbf{c}}}{\partial x_1}(\mathbf{0}), \ldots, \frac{\partial_{\mathcal{C}} I_{\mathbf{c}}}{\partial x_n}(\mathbf{0}) \right)$$

is the $\mathcal{C}$-gradient of $I_{\mathbf{c}}$ at $\mathbf{0}$ (Barceló-Vidal et al., 2011, Definition 13.3.11). Thus

$$\begin{aligned}
I_{\mathbf{c}}(\mathbf{x}) &= I_{\mathbf{c}}(\mathbf{0}) + \sum_{i=1}^{n} \log x_i \frac{\partial_{\mathcal{C}} I_{\mathbf{c}}}{\partial x_i}(\mathbf{0}) \\
&= 0 + \sum_{i=1}^{n} \log x_i \log \frac{c_i}{\mathrm{gm}(\mathbf{c})} \quad \text{(using Equation 5)} \\
&= \sum_{i=1}^{n} \log \left( \frac{1}{n} n x_i \right) \log \frac{c_i}{\mathrm{gm}(\mathbf{c})} \\
&\approx \sum_{i=1}^{n} (-\log n + (n x_i - 1)) \log \frac{c_i}{\mathrm{gm}(\mathbf{c})} \quad \text{(for } n x_i \approx 1\text{)} \\
&= n \sum_{i=1}^{n} x_i \log \frac{c_i}{\mathrm{gm}(\mathbf{c})} \\
&\approx n \sum_{i=1}^{n} x_i \left( \frac{c_i}{\mathrm{gm}(\mathbf{c})} - 1 \right) \quad \text{(for } c_i/\mathrm{gm}(\mathbf{c}) \approx 1\text{)} \\
&= \frac{n}{\mathrm{gm}(\mathbf{c})} \sum_{i=1}^{n} c_i x_i - n.
\end{aligned}$$

Thus the relationship between the Aitchison CTI and the most commonly-used CTI is approximately linear, with slope $n/\mathrm{gm}(\mathbf{c})$ and intercept $-n$, provided the relative abundances (or changes in relative abundances) do not differ too much between species, and the STIs are close to their geometric mean. Note that it would be surprising if the STIs were very different from their geometric mean for a typical community with temperatures measured in Kelvin.
